# Differential Expression Analysis for Longitudinal Single-Cell RNA-Sequencing Studies Using REBEL

**DOI:** 10.64898/2026.05.06.723139

**Authors:** Elizabeth A. Wynn, Kara J. Mould, Brian E. Vestal, Camille M. Moore

## Abstract

Longitudinal scRNA-seq experiments offer a powerful approach for dissecting temporal gene expression dynamics in individual cell types. However, few methods have been developed specifically to address the unique statistical challenges of repeated measures in scRNA-seq data. Here, we introduce a novel method, REBEL (Repeated measures Empirical Bayes differential Expression analysis using Linear mixed models), for analyzing cell type-specific differential expression in repeated measures scRNA-seq experiments. Using simulation studies, we demonstrate that, relative to conventional repeated measures analysis methods and other scRNA-seq approaches, REBEL controls the false discovery rate and exhibits competitive power across a range of simulation scenarios. We further validate REBEL by analyzing a longitudinal scRNA-seq dataset from patients with B-cell lymphoma receiving chimeric antigen receptor (CAR)-T cell therapy. REBEL is implemented as an R package, available at https://github.com/ewynn610/REBEL.

## 1 Introduction

Single-cell RNA-sequencing (scRNA-seq) has emerged as a transformative tool for studying complex biological systems by profiling gene expression at single-cell resolution. A particularly powerful application of scRNA-seq is differential gene expression analysis within specific cell types, an analysis that is not possible with traditional “bulk” RNA-sequencing, where messenger RNA is aggregated across heterogeneous cell populations within a sample.

As sequencing costs continue to decrease, longitudinal and paired studies are becoming increasingly common [Maynard et al., 2020, Wu et al., 2019, Alladina et al., 2023, Kazer et al., 2020]. These designs are well-suited for investigating how gene expression evolves over time within individuals at a cell-type level. However, the analysis of paired or longitudinal scRNA-seq data presents substantial statistical challenges, and rigorously validated methods for this setting remain limited.

A central challenge in longitudinal scRNA-seq differential expression analysis is the hierarchical structure of the data, in which cells are nested within samples, which are themselves nested within subjects. This structure introduces both within-sample and within-subject correlation that must be explicitly modeled. In cross-sectional scRNA-seq studies, within-sample correlation is commonly addressed using the “pseudo-bulk” approach, in which counts for each gene are aggregated across cells within a sample. The resulting data resemble bulk RNA-seq measurements, with one observation per gene per sample, enabling analysis using established bulk RNA-seq methods [Lun and Marioni, 2017, Salcher et al., 2022, Li et al., 2023, Nguyen et al., 2022, Maitra et al., 2023]. In paired or longitudinal designs, however, the correlation between repeated observations from the same subject must also be modeled.

While many traditional bulk RNA-seq tools lack this capability, some methods developed for bulk or single-cell data offer strategies to account for correlation. For instance, limma [Smyth, 2005], initially developed for microarray data and later extended to RNA-seq via the “Voom” transformation [Law et al., 2014], uses linear models and provides the `duplicateCorrelation` function to estimate correlation among repeated measurements [Smyth et al., 2005]. A key limitation of this approach is the assumption that this correlation is constant across all genes, an assumption that is unlikely to hold in practice.

Alternative approaches utilize random-effects models to account for correlated observations. In bulk RNA-seq, methods such as lmerSeq[Vestal et al., 2020] and dream[Hoffman and Roussos, 2021] were developed to handle correlated data using linear mixed models (LMMs). In the single-cell context, related strategies have also been proposed. Dreamlet applies an LMM framework with precision weights to pseudo-bulk data, where the within-sample correlation is addressed throught aggregation and within-subject correlation is modeled via a random subject effect [Hoffman et al., 2023]. Another scRNA-seq method, MAST, employs a two-part hurdle model consisting of a logistic component for expression probability and a Gaussian component for expression magnitude [Finak et al., 2015]. While the MAST package allows random effects to be included in the continuous component, permitting random intercepts for both sample and subject, this usage was not evaluated in the original study.

Although approaches that can be applied to longitudinal scRNA-seq data exist, there has been little systematic benchmarking of differential expression methods for these designs across a range of settings. This is partly because methods such as MAST with random effects and limma with `duplicateCorrelation` are adaptations of existing tools rather than approaches specifically designed for for longitudinal single-cell data. As a result, their relative performance across diverse experimental scenarios remains largely unknown.

Another challenge in longitudinal scRNA-seq analysis is that reliable variance estimation requires adequate sample sizes. In longitudinal bulk RNA-seq studies, both linear and negative binomial mixed models have been shown to yield a high rate of singularities, in which the estimated variance of a random effect collapses toward zero, even in settings where between-subject correlation is explicitly simulated [Vestal et al., 2020, Wynn, 2020]. This issue is likely to extend to scRNA-seq analyses, particularly given the increased noise and sparsity typical of single-cell data. However, current frameworks such as dreamlet and MAST neither report warnings related to singular model fits nor provide estimates random-effect variance to the user.

A well-established strategy for stabilizing parameter estimation in datasets in settings with small sample sizes is empirical Bayes shrinkage, in which information is shared across genes to improve the reliability of variance estimates [Smyth, 2005, Love et al., 2014, Robinson and Oshlack, 2010]. Similar to several bulk RNA-seq methods, dreamlet applies this strategy to residual variance estimates [Hoffman et al., 2023]. Extending this framework, Yu et al. [2019] proposed an approach for repeated measures bulk RNA-seq and microarray studies, that applies empirical Bayes shrinkage to both the residual *and* random-effect variance components. Despite the theoretical advantages of this joint-shrinkage strategy, the absence of accessible software implementations has limited its adoption in scRNA-seq analysis.

To address these challenges, we introduce REBEL (Repeated measures Empirical Bayes differential Expression analysis using Linear mixed models), a method for longitudinal scRNA-seq differential expression analysis. REBEL employs linear mixed models on normalized expression data and implements empirical Bayes shrinkage for both residual and random-effect variance components. The framework supports cell-level (REBEL-cell) and pseudo-bulk (REBEL-PB) differential expression testing. Through simulation studies, we benchmark the performance of REBEL against existing approaches across a range of experimental scenarios, demonstrating its ability to control the false discovery rate while maintaining competitive power. We further illustrate the utility of REBEL by applying it to a longitudinal scRNA-seq dataset from patients with B-cell lymphoma receiving chimeric antigen receptor (CAR)-T cell therapy, revealing a long-term shift in CD8^+^ T cells from a naive/memory precursor state toward a mature effector program post-infusion. REBEL is implemented as an R package and is available at https://github.com/ewynn610/REBEL.

## 2 REBEL Framework

REBEL uses a linear mixed model (LMM) framework applied to variance-stabilized expression data. We implement two modeling approaches: (i) cell-level LMMs, which explicitly model both within-sample and within-subject correlations using random effects, and (ii) a pseudo-bulk approach in which gene expression counts are aggregated within each sample, thereby eliminating within-sample correlation and using a single random effect to account for within-subject correlations. Below we describe the model formulations for each approach.

### 2.1 Linear Mixed Models

Suppose we have a longitudinal scRNA-seq dataset filtered to a single cell type or subpopulation, with gene expression measurements for *m* subjects, each with *n* samples, and an average of *c* cells per sample. For the cell-level approach, we let *Y*_*gijk*_ denote the normalized expression for gene *g* for subject *i*, sample *j*, and cell *k*. The model is specified as:

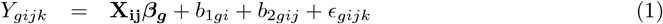

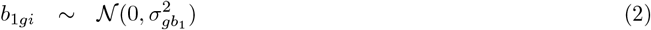

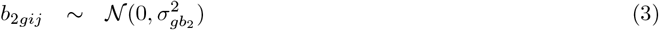

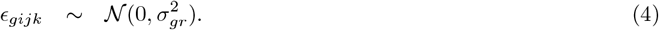

Here, ***β***_***g***_ is a vector of gene-specific fixed-effect coefficients, and **X**_**ij**_ is a corresponding row vector of covariates. The terms *b*_1*gi*_ and *b*_2*gij*_ denote subject- and sample-specific random intercepts, respectively, while *ϵ*_*gijk*_ denotes the residual error, capturing remaining variability. All random effects and residual errors are assumed to be independent and normally distributed with a mean of zero and gene-specific variances.

The pseudo-bulk approach is a reduced form of the cell-level model obtained by aggregating expression across cells within each sample. Let *Y*_*gij*_ denote the normalized pseudo-bulk expression for gene *g*, subject *i*, sample *j*. In this formulation, the fixed effects and subject-level random intercept, *b*_1*gi*_, are retained, the sample-level random intercept is removed, and variability across samples is instead captured by a sample-level residual term, *ϵ*_*gij*_.

For the cell-level approach, normalization and variance stabilization are performed using the `Seurat::SCTransform` function [Hafemeister and Satija, 2019]. For the pseudo-bulk approach, counts are aggregated across cells and transformed using the variance-stabilizing transformation (VST) from the `DESeq2` package [Anders and Huber, 2010, Love et al., 2014].

### 2.2 Empirical Bayes

We extend the empirical Bayes methodology of Smyth [2004], originally developed for differential expression analysis of microarray and bulk RNA-seq data, to the linear mixed model framework utilized in REBEL. Whereas the original framework applies empirical Bayes shrinkage exclusively to residual variance estimates, our approach extends this strategy to both residual and random-effect variance components.

Let 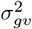 denote a gene-specific variance component for gene *g*, where *v* ∈ {*b*_1_, *b*_2_, *r*} indexes the variance associated with the subject-level random intercept, sample-level random intercept, or residual error, respectively. In the pseudo-bulk approach, the variance components include the subject-level random intercept and residual error (*v* ∈ {*b*_1_, *r*}), whereas the cell-level approach incorporates all three components. Following Smyth [2004], the conditional distribution of the estimator 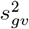 for each variance component is assumed to follow a scaled chi-squared distribution:

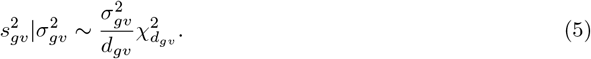

Here, *d*_*gv*_ denotes the associated degrees-of-freedom. These degrees-of-freedom are determined using an adaptation of the between-within degrees-of-freedom approach described by Schluchter and Elashoff [1990] (Supplementary Material section S1).

We place an inverse scaled chi-squared prior on each variance component:

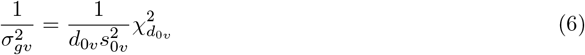

where 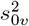 and *d*_0*v*_ are hyperparameters estimated empirically across all genes using the method-of-moments approach of Smyth [2004] (Supplementary Material section S2). Applying Bayes’ theorem yields a closed-form expression for the posterior mean of 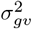 given 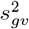:

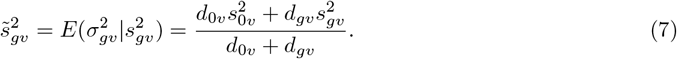

We use 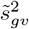 as the empirical Bayes-shrunken estimate of each gene- and component-specific variance parameter.

Hypothesis testing for fixed-effect coefficients or contrasts is performed using *t*-tests, with empirical Bayes-shrunken variance estimates incorporated into the test statistics (Supplementary Material section S3). To conduct inference within the LMM framework, the degrees-of-freedom for the test statistics are estimated using the Kenward–Roger approximation [Kenward and Roger, 1997], substituting the empirical Bayes–shrunken variance estimates into the degree-of-freedom calculations.

## 3 Simulation Study Methods

We conducted a simulation study to evaluate the performance of REBEL in comparison to other potential approaches for longitudinal scRNA-seq differential expression analysis.

### 3.1 Data Generation

Data were simulated using the `rescueSim` R package [Wynn et al., 2025], which generates longitudinal scRNA-seq data using a gamma-Poisson framework that incorporates additional variability at both the sample and subject levels, thereby inducing the correlation structure typical of longitudinal designs. We simulated a two-group study design (control and treatment) with either two or four timepoints per subject (Supplemental Material section S4).

In the two-timepoint design (baseline and follow-up), 20% of genes were simulated to be differentially expressed in the treatment group at the final timepoint, with a log-fold change of ±0.45 (corresponding to approximately a 37% change). In the four-time-point design, these genes followed a linear trajectory over time in the treatment group, reaching the same effect size at the final time point. The number of cells per sample was drawn from a discrete uniform distribution on [0.5*c*, 1.5*c*], where *c* is the average cell count per sample. After simulation, genes expressed in ≤ 10% of cells were filtered from the analysis.

#### 3.1.1 Simulation Scenarios

To evaluate REBEL across a range of experimental conditions, we simulated datasets that varied in the numbers of subjects, timepoints, and average cells per sample, using hyperparameters derived from different biological cell types (Table 1). To assess sensitivity to the strength of within-sample and within-subject correlation, we additionally simulated scenarios in which the location parameters governing between-subject and between-sample variability were shifted by ±*log*(1.3) relative to the baseline (Supplementary Material section S5). For each characteristic, a baseline setting was defined (Table 1), and characteristics was varied independently while holding all other parameters at their baseline values.

**Table 1.** Summary of simulation scenarios. Baseline settings define the reference simulation, and individual data characteristics were varied one at a time while holding all other characteristics fixed at their baseline values.

| Data Characteristic | Baseline Setting | Additional Settings |
| --- | --- | --- |
| Cell type | Recruited Airspace Macrophage | CD4<sup>+</sup> T-cells |
| Subjects per group | 10 | 5, 20 |
| Timepoints per subject | 2 | 4 |
| Average cells per sample | 200 for all groups/timepoints | 50, 100, or 500 for all groups/timepoints<br>500 cells for trt. group/final timepoint,<br>50 cells for other groups/timepoints |
| Subject variability parameter ($\mu_a$) | Estimated from data | $-\log(1.3), +\log(1.3)$ |
| Sample variability parameter ($\mu_b$) | Estimated from data | $-\log(1.3), +\log(1.3)$ |

Simulation parameters were derived from two scRNA-seq datasets representing distinct biological cell types. The first dataset comprised bronchoalveolar lavage (BAL) samples from five healthy adults at baseline and 4-5 days following lipopolysaccharides (LPS) administration (GEO accessions: GSE151928, GSE300946) [Mould et al., 2020]. Cells were integrated using the standard Seurat workflow, clustered via a shared nearest neighbor algorithm, and annotated based on cluster-specific marker genes [Stuart et al., 2019]. Analyses were restricted to recruited airspace macrophages. After filtering out genes with zero expression across all samples, 19,410 genes remained for downstream analysis.

The second dataset comprised blood samples from six children with mild or asymptomatic COVID-19, collected during both the acute and convalescent phases of infection [Khoo et al., 2023]. CD4^+^ T cells were extracted from the processed Seurat dataset available on the GEO database (GEO accession: GSE196456). After filtering out genes with zero expression across all samples, this dataset contained 23,171 genes.

For each simulation scenario, ten datasets were generated. Within each simulation replicate, gene-specific log-fold changes as well as between-subject and between-sample variance parameters were held constant across all scenarios.

### 3.2 Analysis Methods Compared and Implementation

We compared REBEL with both traditional mixed-effects models lacking variance-shrinking adjustments and with existing scRNA-seq and bulk RNA-seq methods that at least partially account for correlation among observations (Table 2). Traditional mixed-effects approaches included negative binomial mixed models (NBMMs) fitted to pseudo-bulk data and linear mixed models (LMMs) applied to variance-stabilized pseudo-bulk and cell-level data. Although we attempted to fit NBMMs at the cell level, the associated computational burden was prohibitive, rendering these analyses infeasible. To mitigate the frequent occurrence of singular fits, we additionally evaluated an alternative strategy wherein singular models were refit after removing the problematic random effect terms (LMM-cell) or by omitting random effects entirely (NBMM-PB and LMM-PB).

**Table 2.** Summary of methods compared.

| Traditional mixed-effect models |  |  |  |  |
| --- | --- | --- | --- | --- |
| Method | Package | Pseudo-bulk/<br>Cell Level | Transformation/<br>Offsets | Implementation Details |
| LMM-cell | `lmerSeq` | Cell | `sctransform` VST | Random effects used for sample/subject. If singular, singular terms removed. Kenward-Roger DF used. |
| LMM-PB | `lmerSeq` | PB | `DESeq2` VST | Subject random effect. Singular models refit as linear model. Kenward-Roger DF used. |
| NBMM-PB | `corrRNASeq` | PB | `DESeq2` offsets | Subject random effect. Singular models refit as negative-binomial linear model. Kenward-Roger DF used. |
| Existing RNA-seq methods |  |  |  |  |
| Method | Package | Pseudo-bulk/<br>Cell Level | Transformation/<br>Offsets | Implementation Details |
| edgeR-PB | `edgeR` | PB | TMM offsets | Default settings. |
| DESeq2-PB | `DESeq2` | PB | `DESeq2` offsets | Default settings. |
| limma-PB | `limma` | PB | Voom + precision weights | Default settings. |
| limma-PB + dupCor | `limma` | PB | Voom + precision weights | `duplicateCorrelation` used with subject as blocking factor. |
| MAST-RE | `MAST` | Cell | $\log(\text{CPM}+1)$ | `glmer` with sample/subject random effects. Cellular detection rate included as fixed effect. |
| dreamlet | `dreamlet` | PB | Voom + weights | Subject random effect and default settings. |
| REBEL-PB | `REBEL` | PB | `DESeq2` VST | Subject random effect. |
| REBEL-cell | `REBEL` | Cell | `sctransform` | Subject/sample random effects. |

Beyond traditional mixed-effects models, we evaluated standard bulk RNA-seq workflows adapted to scRNA-seq data using a pseudo-bulk approach to collapse within-sample correlation (edgeR-PB, DESeq2-PB, and limma-PB). We additionally included specialized methods designed to accommodate both within-sample and within-subject correlation structures, namely limma-PB with duplicate correlation (limma-PB + dupCor), MAST with random effects (MAST-RE), and dreamlet. Additional implementation details for all evaluated methods are provided in Table 2.

All methods were implemented in R version 4.4.1 on a Linux high-performance computing (HPC) cluster. DESeq2-PB, edgeR-PB, and limma-PB, limma-PB + dupCor were executed serially because their runtimes were sufficiently short that parallelization provided no practical benefit. All other methods were run in parallel using 8 cores. For each method, models included group, time, and their interaction, with all variables treated as factors. We evaluated the false discovery rate (FDR) control and power of each method across three specific contrasts: (1) a within-subject test assessing differences between timepoints in the treatment group, (2) a between-subject test assessing differences between groups at the final timepoint, and (3) an interaction test assessing differences in the change over time between groups. Statistical significance was defined using Benjamini-Hochberg adjusted *p*-value threshold of 0.05 for [Benjamini and Hochberg, 1995].

## 4 Methods for Application to Clinical scRNA-seq Data

We demonstrate the practical utility of REBEL through the analysis of a longitudinal scRNA-seq dataset from a study of patients with large B cell lymphoma (LBCL) treated with CD19-directed CAR T cell therapy [Cheloni et al., 2025] (GEO Accession GSE290722). This study employed single-cell transcriptomic profiling of peripheral blood mononuclear cell (PBMC) samples collected at baseline (leukapheresis) and at multiple post-infusion timepoints.

To evaluate the long-term transcriptional response to therapy, we focused on comparisons between baseline and the 6-month follow-up within the CD8^+^ T-cell population. Although the original study characterized subjects by various clinical response outcomes, our preliminary analyses across all methods revealed negligible group-level differences for this cell type at these timepoints. Accordingly, we analyzed subjects as a single cohort to identify overarching longitudinal signatures of CD8^+^ T-cells. We included all samples from these timepoints that had at least 10 CD8^+^ T cells, resulting in a final dataset of 48 samples from 31 subjects with a median of 1,521 cells per sample.

### 4.1 Data Preprocessing and Method Implementation

Prior to analysis, genes expressed in *<* 10% of cells, resulting in a final set of 5,184 genes. We analyzed the data using REBEL alongside the previously described RNA-seq analysis methods. Owing to the large number of cells in the dataset, cell-level methods (REBEL-cell, MAST-RE) were computationally prohibitive on the full dataset and were therefore applied to datasets downsampled to a maximum of 500 cells per sample for samples exceeding this threshold. Pseudo-bulk methods were applied to both the full and downsampled datasets.

For each method, models including fixed effects for timepoint (leukapheresis or 6-month follow-up) and sequencing chip to account for technical batch effects. Where applicable, random effects for sample and/or subject were incorporated. Differential expression was assessed between timepoints for all methods, and resulting *p*-values were adjusted for multiple testing using the Benjamini-Hochberg procedure.

To assess the biological relevance of the differential expression results obtained with REBEL, we performed pathway enrichment analysis on genes identified by REBEL-PB using the `enrichR` package [Kuleshov et al., 2016]. Upregulated and downregulated genes were analyzed separately against the CellMarker 2.0 database [Hu et al., 2023] to characterize shifts in CD8^+^ T-cell transcriptional states. Resulting *p*-values were adjusted for multiple testing using the Benjamini-Hochberg procedure.

## 5 Simulation Results

### 5.0.1 Comparison of REBEL with Empirical Bayes to Standard Mixed Models

We first assessed whether empirical Bayes shrinkage improved performance relative to standard mixed model implementations. Both cell-level and pseudo-bulk mixed models exhibited a substantial proportion of singular fits, in which one or more random effect variance components collapsed to zero, across all simulation scenarios (Fig. 1a, Supplementary Fig. S1). Under the baseline setting, LMM-cell showed the highest rate of singularity (mean = 51.4%), followed by NBMM-PB (38.3%) and LMM-PB (38.3%). Notably, these singular fits occurred despite all genes being simulated with non-zero variance components, indicating that the issue reflects limitations of model estimation rather than an absence of underlying variability.

**Figure 1:**
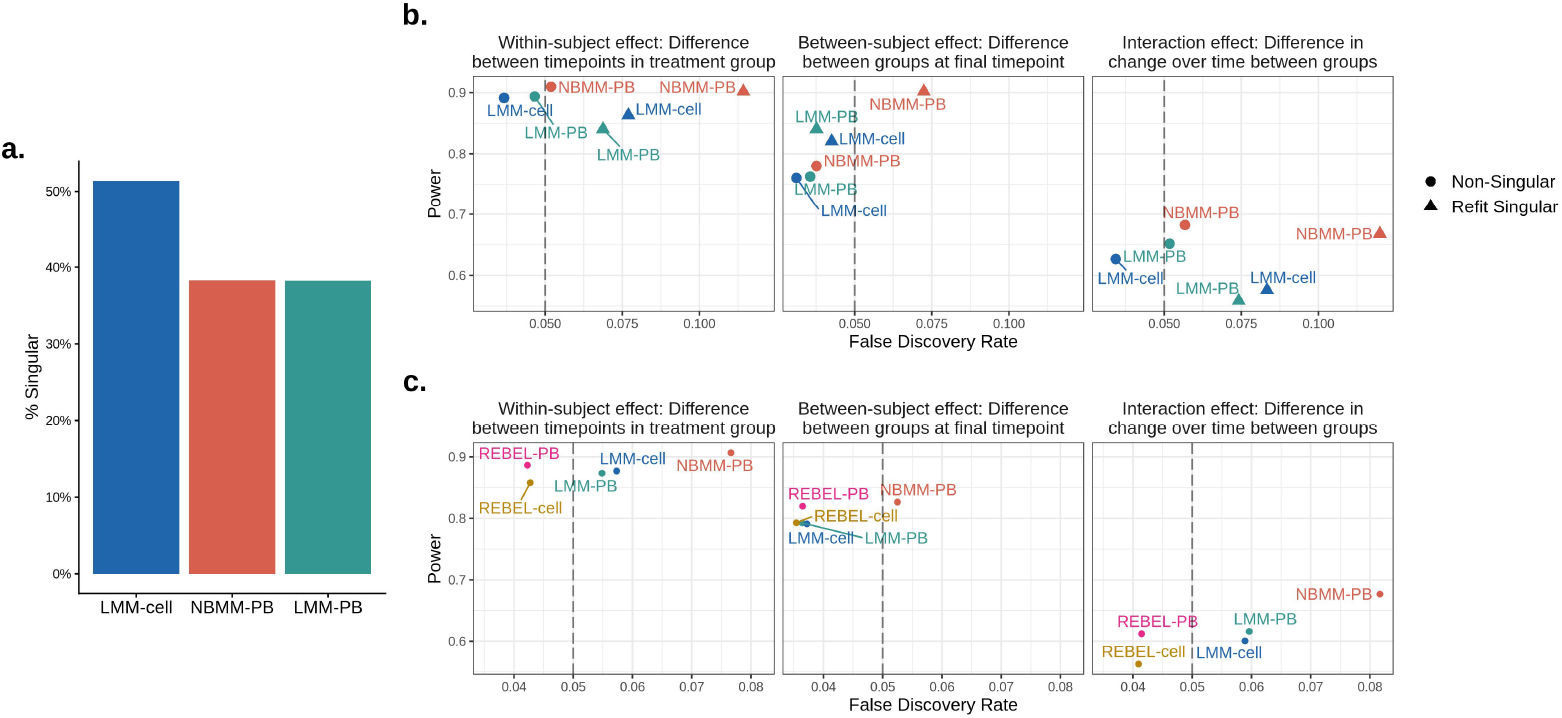
Performance comparison of traditional mixed-effect models and REBEL in longitudinal scRNA-seq simulations. (a) Proportion of singular fits for traditional mixed-effect model implementations under the baseline scenario. (b) Comparison of observed false discovery rate (FDR) and power between non-singular models and singular models refit after removing the zero-variance random effect terms (“reduced” models). (c) Observed FDR versus power for REBEL-PB and REBEL-cell relative to traditional mixed-effect and pseudo-bulk approaches. Results are averaged across ten independent simulations for three contrasts. Statistical significance was defined using a Benjamini-Hochberg adjusted *p*-values threshold of 0.05. Dashed lines denote the nominal 5% false discovery rate.

Singular fits occurred more frequently for genes with lower between-subject variability and higher between-sample variability (Supplementary Fig. S2a). In cell-level models, singularities were more common in the subject random effect and among genes with a higher proportion of zero counts (Supplementary Fig. S2b). Across all tests, models that were singular and subsequently refit after removing random-effect terms with estimated variance equal to zero exhibited increased false discovery rates and reduced power relative to non-singular models. (Fig 1b). These findings suggest that simplification of the random-effects structure in response to singular fits does not recover reliable inference.

When results from non-singular models and singular models refit after removing zero-variance random-effect terms were pooled, LMM-PB, LMM-cell, and NBMM-PB consistently exhibited inflated FDR for the within-subject and interaction tests under the baseline settings. In contrast, REBEL-cell and REBEL-PB successfully maintained proper FDR control while achieving power comparable to the traditional implementations (Fig. 1c). For the between-subject contrast, performance was broadly similar across methods, although NBMM-PB displayed modest FDR inflation. These patterns were consistent across alternative simulation settings (Supplementary Figs. S3-S5).

### 5.1 Comparison of REBEL with Other Single Cell Differential Expression Approaches

We next compared REBEL-PB and REBEL-cell with established single-cell differential analysis frameworks, including MAST-RE and dreamlet, as well as bulk RNA-seq methods adapted for scRNA-seq analysis using a pseudo-bulk strategy (edgeR-PB, DESeq2-PB, limma-PB and limma-PB with duplicate correlation). Among these, edgeR-PB, DESeq2-PB, and limma-PB analyze pseudo-bulk data without explicitly modeling subject-level correlation. Because these three methods exhibited largely comparable performance across scenarios, we focus on limma-PB in the main text for simplicity, with complete results for DESeq2-PB and edgeR-PB provided in the Supplementary Materials (Figures S6-S8).

Under the baseline simulation settings, MAST-RE exhibited substantial FDR inflation across all tests; dreamlet failed to maintain FDR at the nominal level for within-subject and interaction tests, and limma-PB with duplicate correlation showed slightly inflated FDR for the interaction test (Fig. 2a). In contrast, REBEL-PB, REBEL-cell and limma-PB achieved good FDR control, with less than 5% false discoveries, across all tests. Among these three methods, REBEL-PB attained the highest power for within-subject and interaction effects, followed by REBEL-cell and limma-PB. Notably, REBEL-PB achieved similar or higher power than limma-PB with duplicate correlation while also having lower FDR. Although dreamlet and MAST-RE exhibited slightly higher power than REBEL for within-subject and interaction tests, this increase came at the cost of inflated FDR. Though not of primary interest in mixed effects models, FDR and power were broadly comparable across methods for the between-subject test, with the exception of MAST-RE, which had high FDR as noted above.

**Figure 2:**
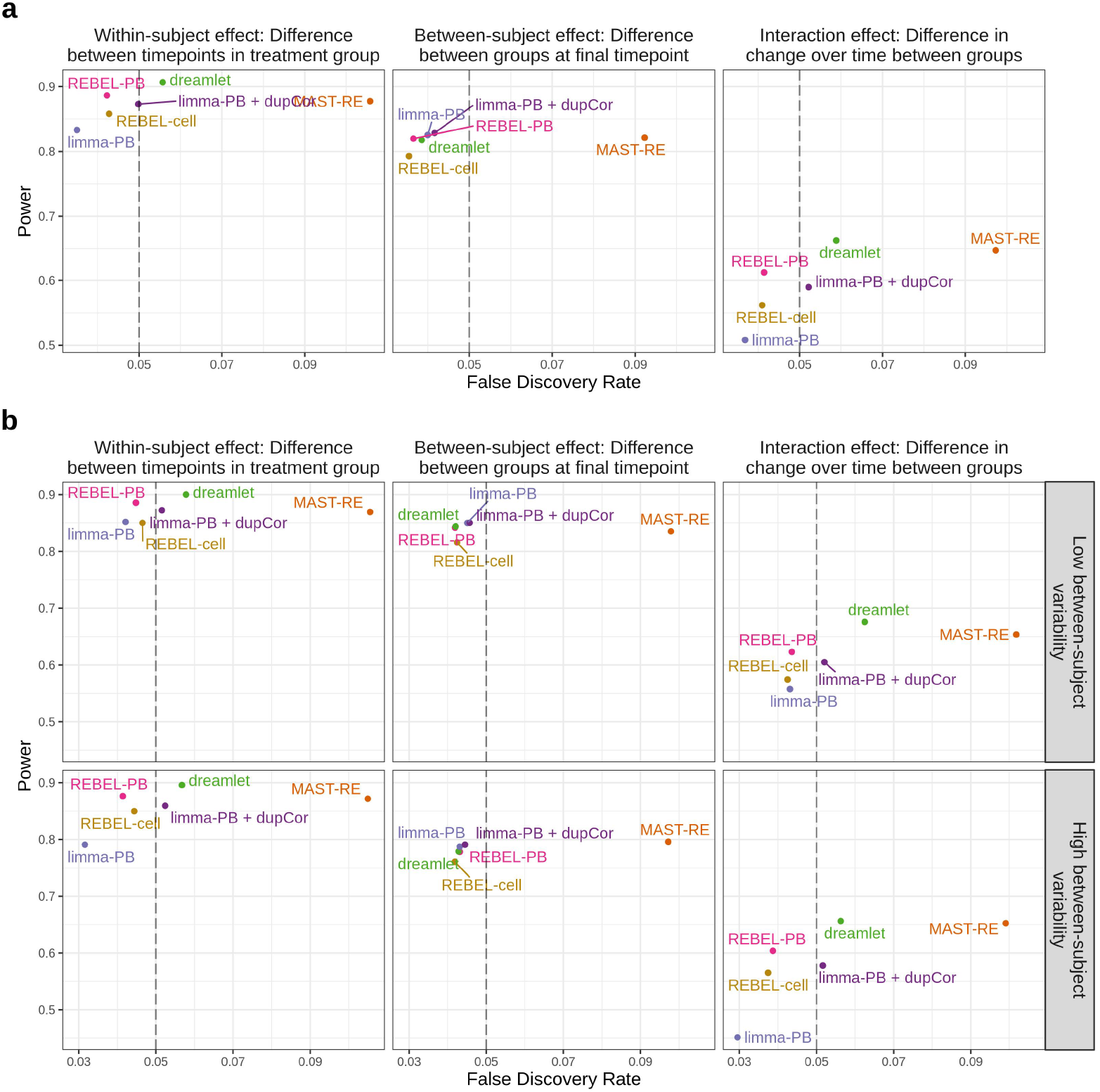
Observed false discovery rate (FDR) versus power for RNA-seq methods that account for correlation under (a) baseline simulation settings and (b) high and low between-subject variability settings. Ten simulated datasets were analyzed for each scenario and the FDR and power were averaged across simulation replicates. Results are shown across the three tests of interest. Results are shown for the three contrasts of interest. Statistical significance was defined using a Benjamini-Hochberg adjusted *p*-value threshold of 0.05. Dashed lines denote the nominal 5% false discovery rate.

#### 5.1.1 Influence of Between-Subject Variability on Method Performance

To assess the impact of within-subject correlation strength influenced method performance, we simulated data with high and low between-subject variability (Fig. 2b). Between-subject tests were largely unaffected across methods and scenarios. For within-subject and interaction tests, the most notable effect of increasing between-subject variability was a reduction in power for limma-PB, which dropped by 7% for the within-subject test and 19% for the interaction test. A modest power reduction with increasing variability was also observed for the interaction test across most other methods (3–5%), though this was far less pronounced. FDR patterns broadly mirrored those observed under baseline conditions, with some methods exhibiting a slight shift toward lower FDR under high between-subject variability relative to low variability for the within-subject and interaction tests. Both REBEL-PB and REBEL-cell maintained well-controlled FDR across both scenarios. When between-sample (rather than between-subject) power reduced for within-subject and interaction tests across all methods (Supplementary Fig. S8a).

#### 5.1.2 Influence of Sample Size on Method Performance

We next examined how variation in the number of subjects, number of timepoints per subject, and number of cells per sample influenced statistical power and FDR. As expected, power increased with the number of subjects, timepoints, and cells per sample (Fig. 3a-b, Supplementary Fig. S8b).

**Figure 3:**
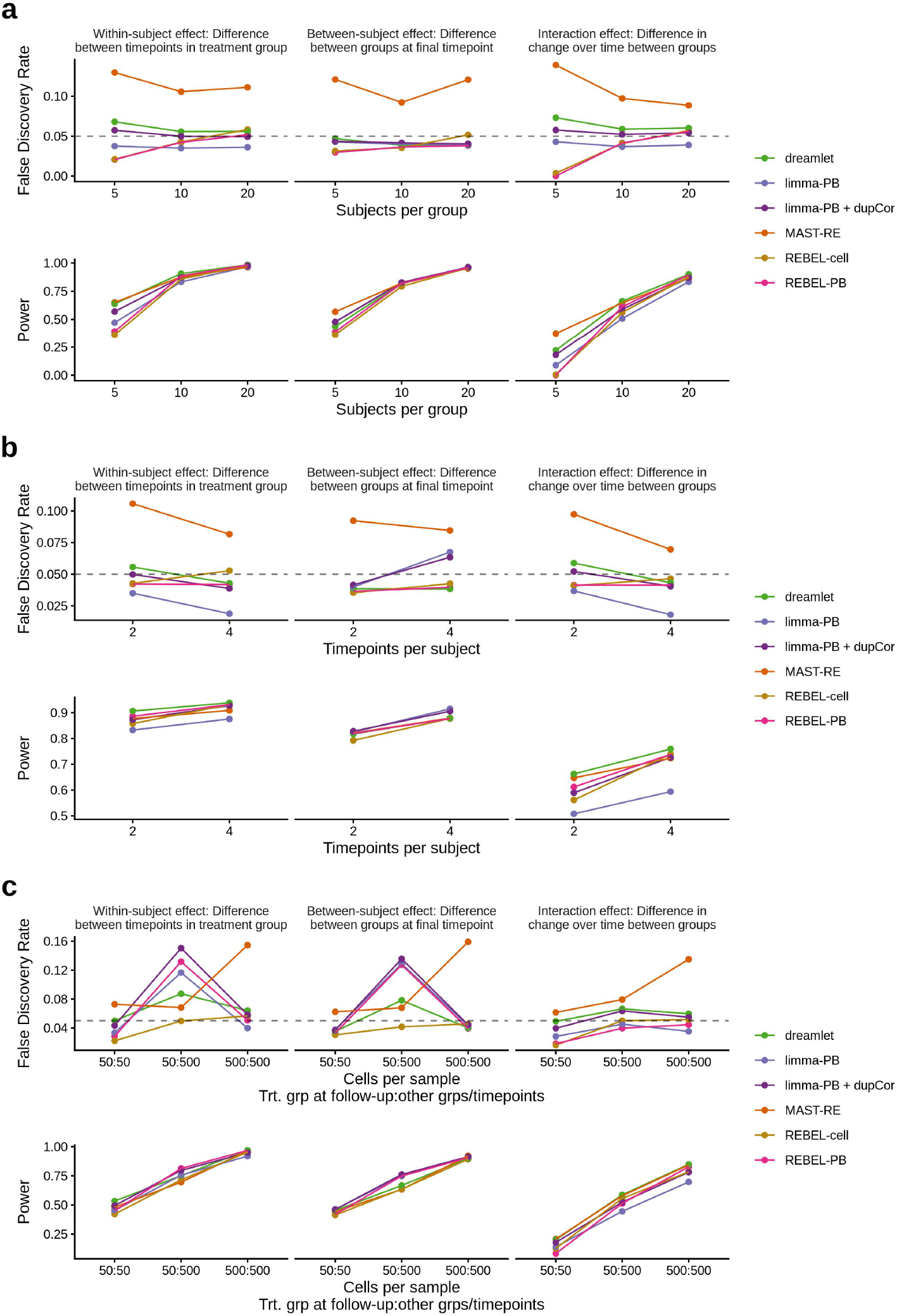
Observed FDR and power for RNA-seq methods that account for correlation across different scenarios of a) number of subjects per group, b) Number of timepoints, and c) average number of cells per group. In c) the x-axis labels contain two numbers with the first number showing the average number of cells in the treatment group at the final timepoint and the second number representing the average number of cells for samples in all other group/timepoint combinations. Results are shown across the three tests of interest. Ten simulated datasets were analyzed for each scenario and the FDR and power were averaged across datasets. A 0.05 threshold was used to determine significance. The dashed lines in the FDR plots represent the nominal rate of 5% false discoveries.

Under the smallest sample size (five subjects per group), all methods exhibited low power across tests, though REBEL-PB and REBEL-cell exhibited the largest reduction (Fig. 3a). In this setting, dreamlet, MAST-RE, and limma-PB with duplicate correlation additionally had inflated FDR for the within-subject and interaction tests, whereas REBEL remained conservative. With increasing sample size, the FDR for REBEL-PB and REBEL-cell increased, and the FDR inflation for dreamlet and limma-PB with duplicate correlation decreased to near nominal levels. MAST continued to show FDR inflation across all tests even at the largest sample size, while limma-PB was conservative.

With four timepoints per subject, FDR remained conservative for REBEL-PB and increased to near nominal levels for REBEL-cell (Fig. 3b). For all other methods, FDR decreased for within-subject and interaction tests. Interestingly, limma-PB and limma-PB with duplicate correlation showed elevated FDR for the between-subject test.

Across varying numbers of cells per sample (50, 100, 200, and 500), differences in method performance were largely consistent with the baseline, though MAST-RE showed increasing FDR inflation at higher cell counts (Supp Fig. S3b). Interestingly, in the unbalanced cell count scenario, where the follow-up timepoint in the treatment group had substantially fewer cells per sample than all other group/timepoint combinations, pseudo-bulk methods showed inflated FDR for between-subject and within-subject tests, while REBEL-cell maintained well-controlled FDR across all tests (Fig. 3c). This may represent a key advantage of the cell-level approach when cell counts are highly unbalanced.

#### 5.1.3 Simulations on a Different Cell Type

Simulations using a different dataset and cell type (CD4^+^ T cells) yielded broadly similar results to the macrophage baseline, though power and FDR control were generally reduced across methods. This suggests that cell type-specific characteristics influence method performance even when experimental design parameters such as the number of subjects and cells are held constant.

## 6 Application to Clinical scRNA-seq Data

We applied REBEL, alongside existing approaches, to CD8^+^ T cells from a longitudinal scRNA-seq dataset of patients undergoing CD19-directed CAR T cell therapy [Cheloni et al., 2025]. To investigate the long-term transcriptional remodeling of the native T-cell population, we performed a within-subject test comparing gene expression at leukapheresis (baseline) versus 6-months post-infusion.

### 6.1 Comparative Performance of Analysis Methods

Due to the large number of cells in the dataset, cell-level methods (MAST-RE and REBEL-cell) were implemented using data downsampled to a maximum of 500 cells per sample to ensure computational feasibility. All other methods were applied to the full dataset. To assess whether this downsampling influenced our findings, methods run on the full dataset were also applied to the downsampled data. The results were broadly consistent, with the downsampled analysis yielding a similar core set of differentially expressed genes (DEGs) with a marginal reduction in the total count of DEGs (Supplementary Fig. S9).

The number of differentially expressed genes (DEGs) detected across methods broadly reflected performance patterns observed in simulations (Fig 4a). MAST-RE identified the most DEGs by far (2,925), consistent with its substantial FDR inflation across all simulation scenarios, particularly with a large number of cells per sample (Supplementary Fig. S8b). Dreamlet also identified notably more DEGs than other methods (417), aligning with simulation patterns where its higher power was accompanied by FDR above the nominal threshold for the within-subject effect across most scenarios.

**Figure 4:**
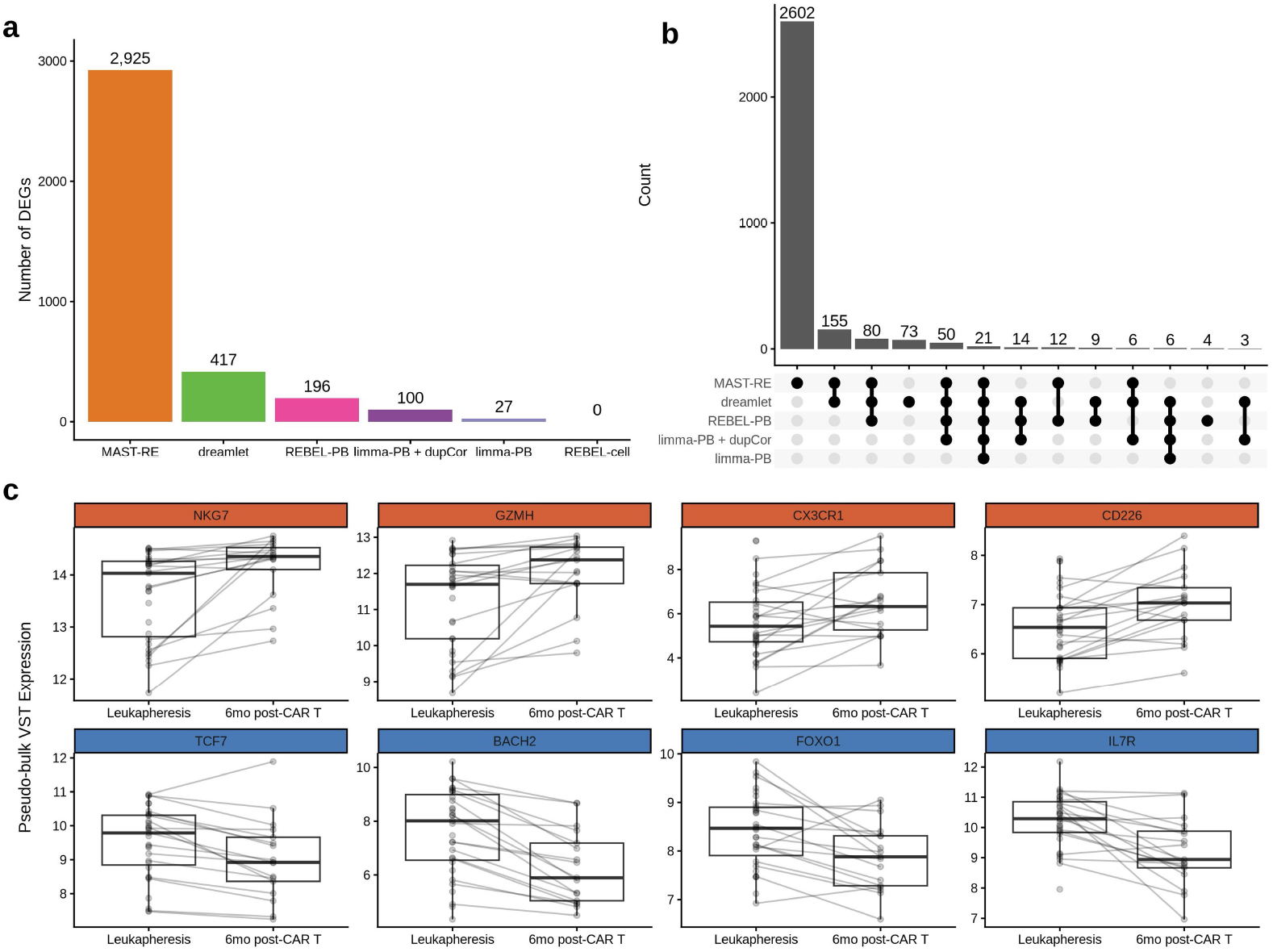
Longitudinal differential expression analysis of CD8^+^ T cells following CAR T-cell therapy (a) Number of differentially expressed genes (DEGs) identified by REBEL-PB and competing longitudinal methods. (b) UpSet plot displaying the intersection of DEGs across all tested methods, illustrating the unique and shared gene sets identified. (c) Paired expression of key cytotoxic and naive markers at leukapheresis (Baseline) and six months post-infusion. Each point represents the variance-stabilizing transformed (VST) expression of a single patient’s native CD8^+^ T cells, with lines connecting paired samples. Gene titles highlighted in red (NKG7, GZMH, CX3CR1, CD226) indicate significant upregulation at six months post-infusion, while those in blue (TCF7, BACH2, FOXO1, IL7R) indicate significant downregulation.

REBEL-PB identified 196 DEGs, while limma-PB with duplicate correlation identified 100. Standard limma-PB found only 27 DEGs, a result likely attributable to the dramatic power loss observed in our simulations under high between-subject variability. Notably, the subject-level ICC values for this dataset, calculated using the REBEL-PB random effects, were comparable to those in the high between-subject variability simulation scenario, with this dataset having an average ICC of 0.26 and the simulations having an average ICC of 0.256. Finally, REBEL-cell, which had among the lowest power across scenarios, found no DEGs.

We next examined the overlap in DEGs identified across the different methods (Fig. 4b). The DEGs identified by limma-PB and limma-PB with duplicate correlation were largely subsets of those identified by REBEL-PB. Specifically, all 27 limma-PB DEGs and 91 of 100 limma-PB with duplicate correlation DEGs were also identified by REBEL-PB, suggesting these methods are recovering a consistent core set of differentially expressed genes. While MAST-RE and dreamlet recovered the majority of this core set as well, with only 4 REBEL-PB DEGs found by neither method, they additionally identified a vast number of genes found by no other method. Specifically, 2,602 genes were exclusive to MAST-RE, 73 were exclusive to dreamlet, and 155 shared by those two methods. This high frequency of exclusive DEGs is consistent with the inflated FDR observed for these methods in our simulations, suggesting that many of these additional findings may represent false discoveries.

### 6.2 Biological Characterization of REBEL-PB Results

We examined the results from REBEL-PB more closely to assess whether the detected DEGs reflected biologically coherent, long-term transcriptional changes following CAR T cell therapy. First, we performed gene set enrichment analysis using the CellMarker 2.0 database [Hu et al., 2023] to characterize the biological signatures of REBEL-PB DEGs (Supplementary Table S1). Upregulated genes were significantly enriched for transcriptional programs associated with effector memory CD8^+^ T-cell function (Top term: Effector CD8+ Memory T (Tem) Cell Peripheral Blood Human, *p* = 9.12 × 10^−19^, 13/77 overlapping genes), while downregulated genes were enriched for naive and memory precursor CD8^+^ T-cell signatures (Top term: Naive CD8+ T Cell Peripheral Blood Human, *p* = 7 × 10^−8^, 11/97 overlapping genes). This pattern indicates that the patients’ native T cells moved away from a resting state and transitioned into a more mature, active state.

In support of this shift, REBEL-PB identified the upregulation of several key markers of cytotoxic effector function and terminal differentiation, including NKG7, GZMH, CX3CR1, and CD226 [Turiello et al., 2025, Böttcher et al., 2015, Vo et al., 2016] (Fig. 4c). Conversely, downregulated genes included transcription factors and receptors associated with T cell quiescence and long-term maintenance, including TCF7, BACH2, FOXO1, and IL7R [Delpoux et al., 2017, Conti et al., 2026], further supporting a broad transition away from a resting, stem-like state toward an active effector function.

Taken together, these results are consistent with a shift in the CD8^+^ T-cell transcriptional program from a naive/memory precursor state at leukapheresis toward a mature effector program by 6-months post-infusion. This reflects a durable, longer-term remodeling of the native CD8^+^ T-cell compartment following CAR T-cell therapy. Such a transition toward a more cytotoxic effector state is consistent with previous reports of stable host immune remodeling following CAR T-cell infusion [Chen et al., 2020].

## 7 Discussion

The increasing prevalence of longitudinal scRNA-seq studies necessitates statistical frameworks that can accurately model the complex, hierarchical structure of longitudinal scRNA-seq designs. In this paper, we introduced a novel method, REBEL, for analyzing longitudinal scRNA-seq data. Our method uses an LMM framework on transformed data with residual and random effect variances estimated using an empirical Bayes process, allowing the sharing of information across genes. REBEL can be implemented on both pseudo-bulk and cell-level data.

A central challenge in analyses of longitudinal scRNA-seq data is the instability of variance component estimation. We demonstrated that standard traditional mixed model approaches frequently produce singular fits, even when the random effect variances are simulated to be strictly positive. While common practice suggests simplifying these models by removing the “offending” random effect, our results show that this leads to inflated FDR and reduced power. By utilizing empirical Bayes shrinkage to stabilize these estimates, REBEL effectively “rescues” these models, borrowing information across the transcriptome to ensure that subject-level and sample-level correlations are accounted for even when individual gene data is sparse.

Through extensive simulations, we also evaluated the performance of REBEL against a broad range of established methodologies that partially or fully attempt to account for data correlation. Our results underscore that ignoring subject-level information leads to a substantial loss in statistical power, a problem that is exacerbated as between-subject variability increases. Notably, this loss of power persists even when utilizing pseudo-bulk approaches on sample-level data; when subject-level correlations were neglected, standard frameworks such as limma-PB, edgeR-PB, and DESeq2-PB struggled to maintain sensitivity. Even when employing limma’s duplicate correlation framework, the method often yielded lower or comparable power while exhibiting higher False Discovery Rates (FDR) than REBEL-PB across numerous scenarios.

Similarly, existing methods designed for scRNA-seq data exhibited notable performance limitations across our simulation scenarios. MAST-RE had severely inflated FDR in nearly scenarios, while dreamlet maintained high power but showed inflated FDR. Relative to other methods, REBEL-PB effectively controlled FDR while maintaining comparable power across diverse experimental conditions.

We observed some differences in performance between the two implementations of our method. Across simulations, REBEL-cell generally exhibited less power than REBEL-PB. However, in the scenario characterized by imbalance in the average number of cells between groups and timepoints, REBEL-cell offered a unique advantage over pseudo-bulk methods. While pseudo-bulk methods exhibited inflated FDR in both within-subject and between-subject tests under these conditions, REBEL-cell maintained robust error control. This is a highly relevant consideration in clinical research, where disease progression, exposures, or treatments may significantly deplete or enhance specific cell populations, leading to systematic imbalances in cell counts across timepoints or groups. In such cases, the cell-level model provides a necessary tool for extracting insights from imbalanced datasets. Consequently, by providing both pseudo-bulk and cell-level options within the REBEL framework, we ensure that researchers can select the model most suited to the specific constraints and biological characteristics of their data.

To evaluate the utility of our framework in a real-world setting, we applied REBEL and competing methods to a clinical longitudinal scRNA-seq dataset of CD8^+^ T cells. The results across methods generally mirrored our simulation findings regarding sensitivity and power. Specifically, MAST-RE and dreamlet identified the highest number of DEGs, while REBEL-PB and limma with duplicate correlation fell into a middle tier of sensitivity. In contrast, limma-PB identified very few DEGs, and REBEL-cell identified none.

The biological insights provided by REBEL-PB demonstrate its ability to extract meaningful signals from complex clinical data. Our findings revealed a significant enrichment for cytotoxic effector programs in genes upregulated at six months post-infusion, contrasted by the downregulation of markers associated with T-cell quiescence and naive states. Ultimately, the ability of REBEL-PB to identify these biologically coherent signatures while maintaining the balanced error control observed in our simulations highlights its value as a rigorous tool for longitudinal scRNA-seq analysis.

There are several limitations to our method and study design that highlight important considerations for researchers and future areas of investigation. First, our results demonstrate that method performance in longitudinal scRNA-seq analysis is somewhat context-specific, with no single framework proving universally superior. While REBEL maintained robust FDR control across most scenarios, it exhibited substantially reduced power in cases with small sample sizes, suggesting it may be less effective for studies with limited cohorts. Furthermore, due to the complexity of scRNA-seq data, our simulations focused on single-factor perturbations from baseline settings and we did not explore the interactions between different factors. Future research characterizing method performance across a broader combination of data characteristics will be essential for developing guidelines on how to select the most appropriate statistical approach for a given dataset.

There are also some practical constraints to the REBEL framework. Like other cell-level methods, REBEL-cell is significantly more resource-intensive than pseudo-bulk approaches. In large clinical datasets, including the clinical example used in this paper, this may necessitate downsampling of cells. More rigorous study is needed to determine the downstream effects and best practices for downsampling approaches. Finally, while our current random-intercept model successfully accounts for primary hierarchical correlations, it does not incorporate random slopes or auto-correlation structures. Expanding REBEL to include these features will be an important next step as scRNA-seq experiments continue to explore increasingly complex temporal dynamics.

The REBEL method provides a statistically rigorous yet flexible framework for the analysis of longitudinal scRNA-seq data, addressing the critical challenge of variance instability through an empirical Bayes approach. By offering both pseudo-bulk and cell-level implementations, REBEL enables researchers to maintain robust error control and biological sensitivity across a diverse range of experimental designs and cell type populations. REBEL is easily implemented using our open source R package hosted on GitHub at https://github.com/ewynn610/REBEL.

