## Supplemental Material for "Differential Expression Analysis for Longitudinal Single-Cell RNA-Sequencing Studies Using REBEL"

#### S1 Degrees of Freedom for Empirical Bayes Variance Estimates

Within the empirical Bayes procedure, the degrees of freedom associated with each variance estimates, $d_{gv}$, must be specified. Smyth [2004] showed that, in a linear model framework, the residual degrees of freedom provide an appropriate choice for modeling the sampling distribution of the residual variance estimator. In the LMM framework used by REBEL, specifying degrees of freedom is more nuanced, as multiple variance components are present and analytical expressions for degrees of freedom are generally unavailable and must be approximated.

We calculate $d_{gv}$ using an approach based on the between-within degrees-of-freedom decomposition commonly employed for hypothesis testing in LMMs. This approach partitions the residual degrees of freedom into between-subject and within-subject components. The between-subject degrees of freedom are used for inference on effects that vary across subjects, whereas the within-subject degrees of freedom are used for effects that vary within subjects. This decomposition aligns naturally with the variance components in our pseudo-bulk model, where the subject-level random effect variance captures variability between subjects and the residual variance captures variability within subjects.

Accordingly, for the subject-level random effect variance, we use the between-subject degrees of freedom, defined as the number of subjects minus the number of fixed-effect covariates that vary between subjects.

For the residual variance, we use the within-subject degrees of freedom, defined as the total residual degrees of freedom minus the between-subject degrees of freedom.

For the cell-level model, we further decompose the residual degrees of freedom into between-subject and between-sample components. The between-subject degrees of freedom are calculated as described above. The between-sample degrees of freedom are obtained by subtracting the number of fixed-effect covariates that vary between samples from the total number of samples. These between-subject and between-sample degrees of freedom are used for the subject- and sample-level random effect variance components, respectively. For the residual variance, we use the between-sample degrees of freedom.

### S2 Prior Parameter Estimation

We estimate the prior parameters $s_{0v}$ and $d_{0v}$ using the empirical Bayes procedure described by Smyth [2004]. Briefly, the theoretical first and second moments of $log(s_{gv}^2)$, which are functions of on $s_{0v}$ and $d_{0v}$, are equated to their empirical counterparts across genes. The resulting equations are then solved to obtain method-of moments estimates for the prior parameters.

Because this procedure operates on the logarithmic scale, estimation of prior parameters for random-effect variance components is not possible for singular model fits in which the estimated random-effect variance is zero. Such singular fits commonly arise spuriously in settings with sparse data or limited sample sizes. Therefore, when estimating the prior parameters for random-effect variance components, genes with singular model fits are excluded from the estimation procedure. Importantly, random-effect variance estimates from singular models are retained and incorporated in the subsequent computation of posterior variance estimates.

### S3 Hypothesis Testing

Hypothesis testing of model coefficients or contrasts is performed using empirical Bayes-shrunken variance estimates to compute standard errors and degrees of freedom. Let $\mathbf{Z}$ denote the variance-covariance matrix of the random effects for gene $g$, constructed using empirical Bayes-shrunken variance estimates. In the pseudo-bulk approach, a single random effect is included and thus $\mathbf{G}_g = \tilde{s}_{gb_1}^2$. In the cell level approach, subject- and sample-level random effects are modeled, yielding

$$\mathbf{G}_g = \begin{bmatrix} \tilde{s}_{gb_1}^2 & 0 \\ 0 & \tilde{s}_{gb_2}^2 \end{bmatrix} \tag{1}$$

The, marginal covariance matrix of the response vector $\mathbf{Y}_g$,is then given by

$$\mathbf{V_g} = \mathbf{ZGZ}^T + \tilde{s}_{gr}^2, \tag{2}$$

where $\tilde{s}_{gr}^2$ denotes the empirical Bayes–shrunken residual variance.

Let $\mathbf{X_g}$ denote the fixed effect design matrix. The covariance matrix of the estimated the fixed-effect coefficients $\boldsymbol{\beta}_g$ is computed as

$$\mathbf{V}(\boldsymbol{\beta}_\mathbf{g}) = (\mathbf{X^T V(Y_g)^{-1} X})^{-1} \tag{3}$$

To test the null hypothesis $H_0 : \mathbf{L}\boldsymbol{\beta}$ for a contrast vector $\mathbf{L}$, we construct a $t$ t-test statistic of the form

$$t = \frac{\mathbf{L}\boldsymbol{\beta}}{\sqrt{\mathbf{LV}(\boldsymbol{\beta}_\mathbf{g})}} \tag{4}$$

which is assumed to follow a $t$ distribution with gene-specific degrees of freedom $d_g$. P-values are computed for each gene and adjusted for multiple testing using the Benjamini-Hochberg procedure [Benjamini and Hochberg, 1995].

##### S3.1 Kenward-Roger Degrees of Freedom and Covariance Adjustment

Degrees of freedom for hypothesis testing are estimated using the Kenward-Roger approximation, which provides a small-sample correction for inference in linear mixed models [Kenward and Roger, 1997]. This method is based on a moment-matching strategy related to the Satterthwaite approximation [Satterthwaite, 1941, 1946]. In addition to estimating degrees of freedom, we use the Kenward-Roger bias-adjusted estimator of the fixed-effect covariance matrix when computing the test statistic in Equation 4). Both the degrees of freedom and the bias-adjusted covariance matrix are computed using the empirical Bayes-shrunken variance component estimates.

#### S4 Data Simulation

Let $Y_{gijk}$ denote the raw count value for gene $g$, subject $i$, sample $j$, and cell $k$. Let $L_{ijk}$ denote the expected library size for subject $i$, sample $j$ and cell $k$ drawn from a log-normal distribution with parameters estimated using empirical data. Then,

$$Y \sim Poisson(x^*_{gijk}) \tag{5}$$

$$x^*_{gijk} = L_{ijk} \times \frac{x_{gijk}}{\sum_g x_{gijk}} \tag{6}$$

$$x_{gijk} \sim Gamma\Big(shape = \frac{1}{\phi_g}, scale = \phi_g \mu_{gij} z_{gij}\Big) \tag{7}$$

$$log(z_{gij}) = \beta_{g1} X_{1i} + \beta_{g2} X_{2ij} + \beta_{g3} X_{1i} X_{2ij} \tag{8}$$

Here, $\mu_{gij}$ represents the mean expression level of gene $g$ for sample $j$ from subject $i$. Sample-level values of $\mu_{gij}$ are derived from empirical data and simulated to induce stronger correlation among samples from the same subject than among samples from different subjects. The dispersion parameter $\frac{1}{\phi_g}$ is gene specific and is also estimated from empirical data. The scaling factor $z_{gij}$ is used to introduce systematic differential expression across experimental conditions.

The indicator variable, $X_{1i}$ denotes whether subject $i$ belongs to the treatment group, and $X_{2ij}$ encodes the time point associated with sample $j$ from subject $i$, taking values in $0, 1, ...t$ where $t + 1$ is the number of time points. The value $\beta_{g1}$ represents the log-fold change between treatment and control groups at the baseline, $\beta_{g2}$ represents the log-fold change across time points in the control group, and $\beta_{g3}$ captures the group-by-time interaction effect.

Differential expression is simulated by setting $\beta_{g1} = \beta_{g2} = 0$ for all genes across all genes and assigning $\beta_{g3} = \frac{0.45}{t}$ in 20% of genes. Under this specification, differentially expressed genes exhibit a total log-fold change of 0.45 (approximately a 37% difference) between the first and final time points in the treatment group, or between treatment and control groups at the final time point.

### S5 Modulation of Simulation Subject- and Sample-Level Variability

The longitudinal scRNA-seq data were simulated using the `rescueSim` package, which introduces hierarchical deviations around a global mean expression level $\mu_g$ for each gene. Specifically, the mean expression for gene $g$ in subject $i$ and sample $j$ ($\mu_{gij}$) is defined as:

$$\mu_{gij} = \mu_g a_{gi} b_{gij} \tag{9}$$

where $a_{gi}$ and $b_{gij}$ are multiplicative factors representing subject- and sample-specific deviations, respectively. These deviation factors are drawn from Dirichlet distributions with gene-specific concentration parameters determined by their corresponding variability parameters, $v_{ag}$ and $v_{bg}$. To model heterogeneity in the variability across the the transcriptome, the gene-level variability parameters are themselves drawn from log-normal distributions:

$$v_{ag} \sim \text{LNorm}(\mu_a, \sigma_a^2) \tag{10}$$

$$v_{bg} \sim \text{LNorm}(\mu_b, \sigma_b^2) \tag{11}$$

The hyperparameters $\mu_a$ and $\mu_b$ represent the log-scale mean subject- and sample-level variability, respectively. To assess sensitivity to the strength of within-subject and within-sample correlation, we modulated these parameters by shifting their baseline values by $\pm 1.3$, while holding all other simulation settings fixed.

#### S6 Supplemental Figures

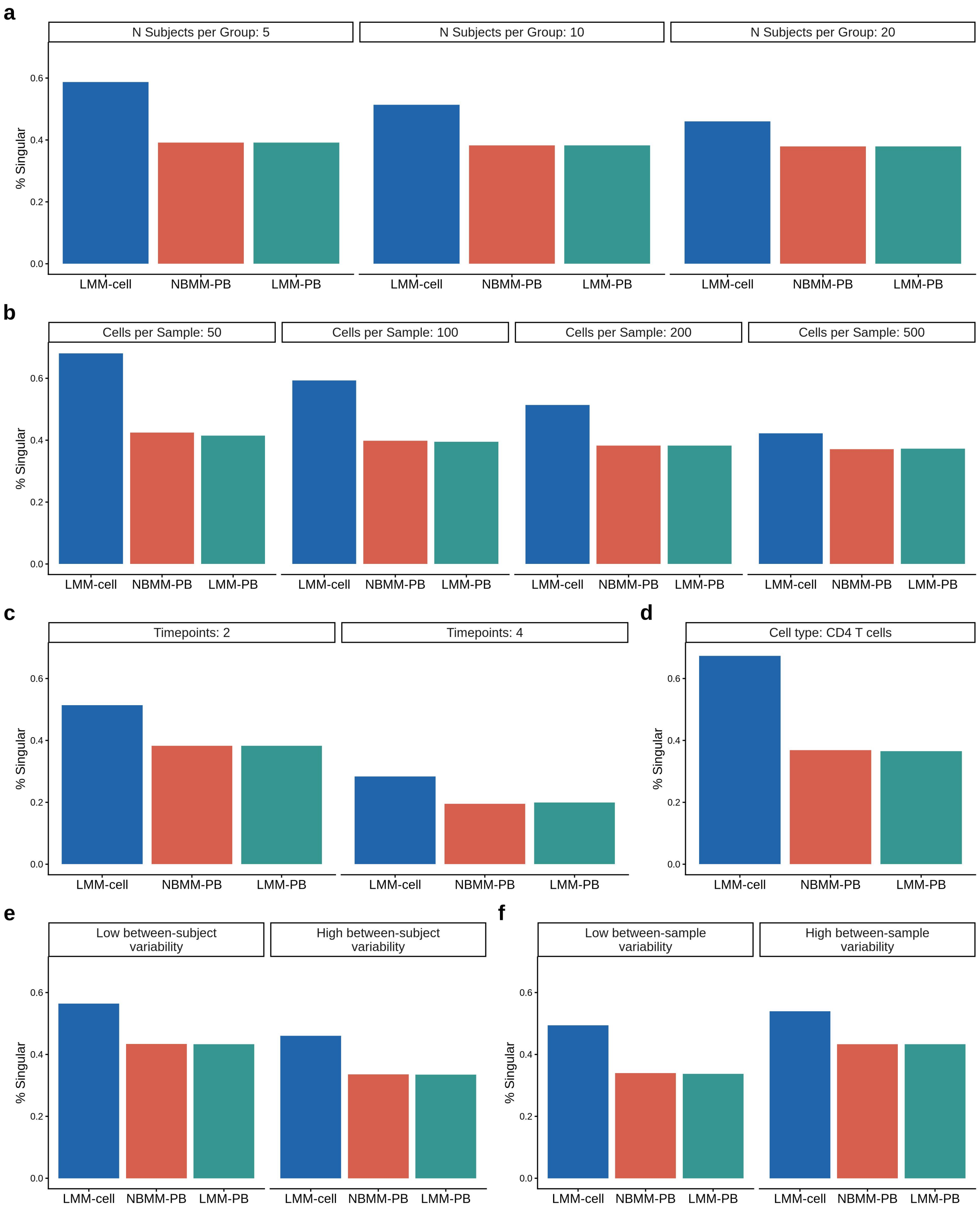


Figure S1: Proportion of singular fits for traditional mixed-effects model implementations across simulation scenarios. Panels show singularity rates as a function of varying (a) number of subjects per group, (b) average number of cells per sample, (c) number of timepoints per subject, (d) simulated cell type ($CD4^+$ T cells), (e) increased or decreased between-subject variability, and (f) increased or decreased between-sample variability. For each scenario, only the indicated characteristic was varied while all other simulation parameters were held at their baseline values. Results are averaged across ten independent simulation replicates.

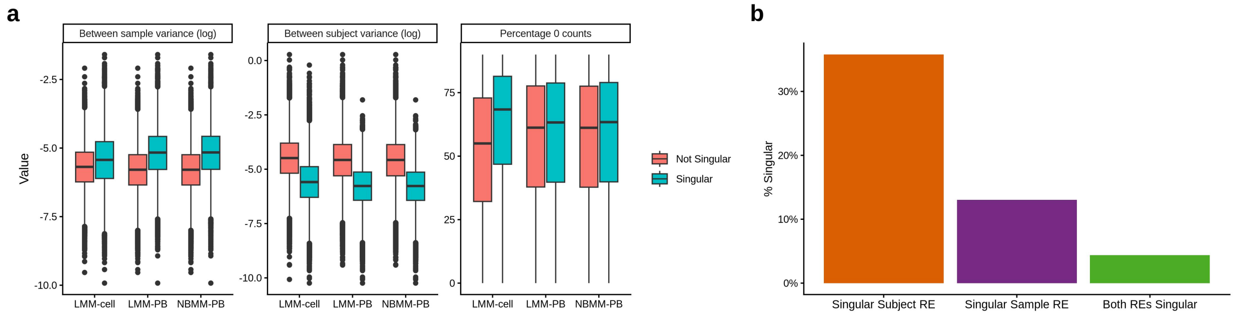


Figure S2: Characteristics and structure of singular fits in traditional mixed-effects models. (a) Distributions of gene-level characteristics stratified by whether models produced singular or non-singular fits, including between-sample variance simulation parameter, between-subject variance simulation parameter, and percentage of zero counts. Results are shown for LMM-cell, LMM-PB, and NBMM-PB implementations. (b) Proportion of singular fits in LMM-cell models attributable to singularity in the subject-level random effect, the sample-level random effect, or both random effects simultaneously.

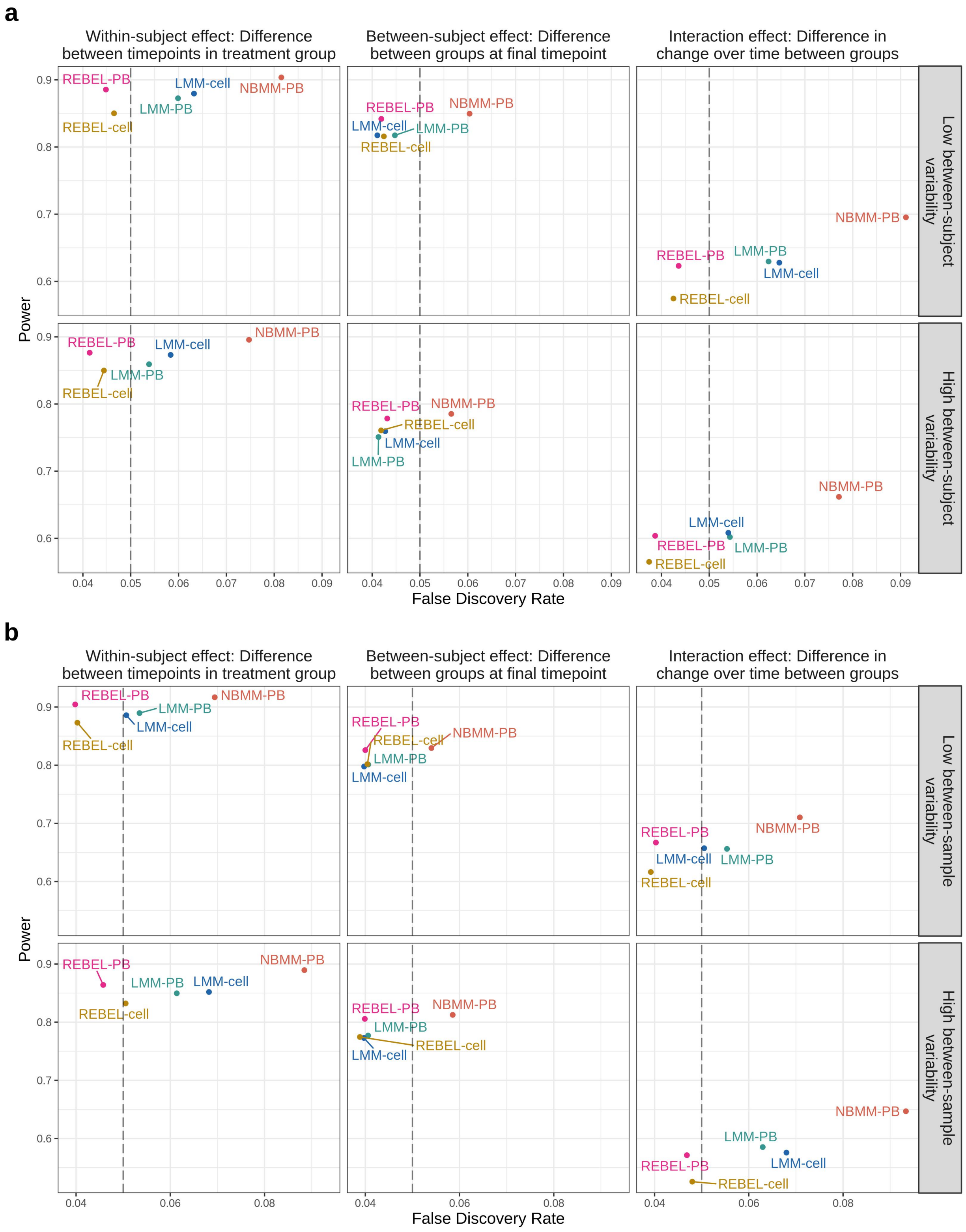


Figure S3: Observed false discovery rate (FDR) versus power for traditional mixed-effect models and REBEL implementations under (a) high and low between-subject variability settings and (b) high and low between-sample variability settings. Ten simulated datasets were analyzed for each scenario and the FDR and power were averaged across datasets. Results are shown for the three contrasts of interest. Statistical significance was defined using a Benjamini-Hochberg adjusted $p$-value threshold of 0.05. Dashed lines denote the nominal 5% false discovery rate.

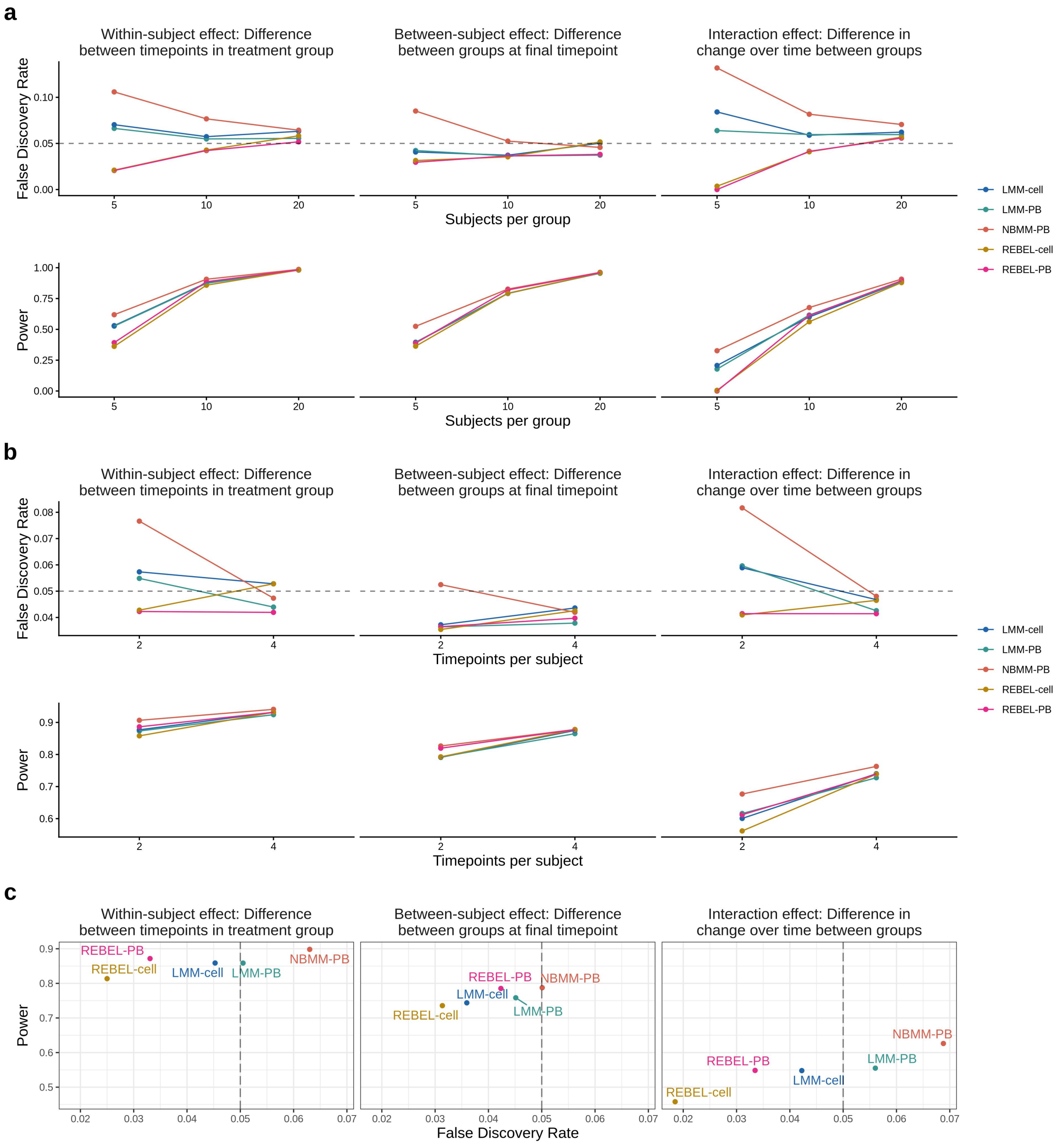


Figure S4: Observed false discovery rate (FDR) and power for traditional mixed-effect models and REBEL implementations across alternative simulation scenarios. Results are shown as a function of varying (a) number of subjects per group, and (b) number of timepoints per subject. Panel (c) displays observed FDR versus power for the CD4$^+$ T-cell simulation scenario. Results are shown for the three contrasts of interest. Statistical significance was defined using a Benjamini-Hochberg adjusted $p$-value threshold of 0.05. Dashed lines in the FDR plots denote the nominal 5% false discovery rate.

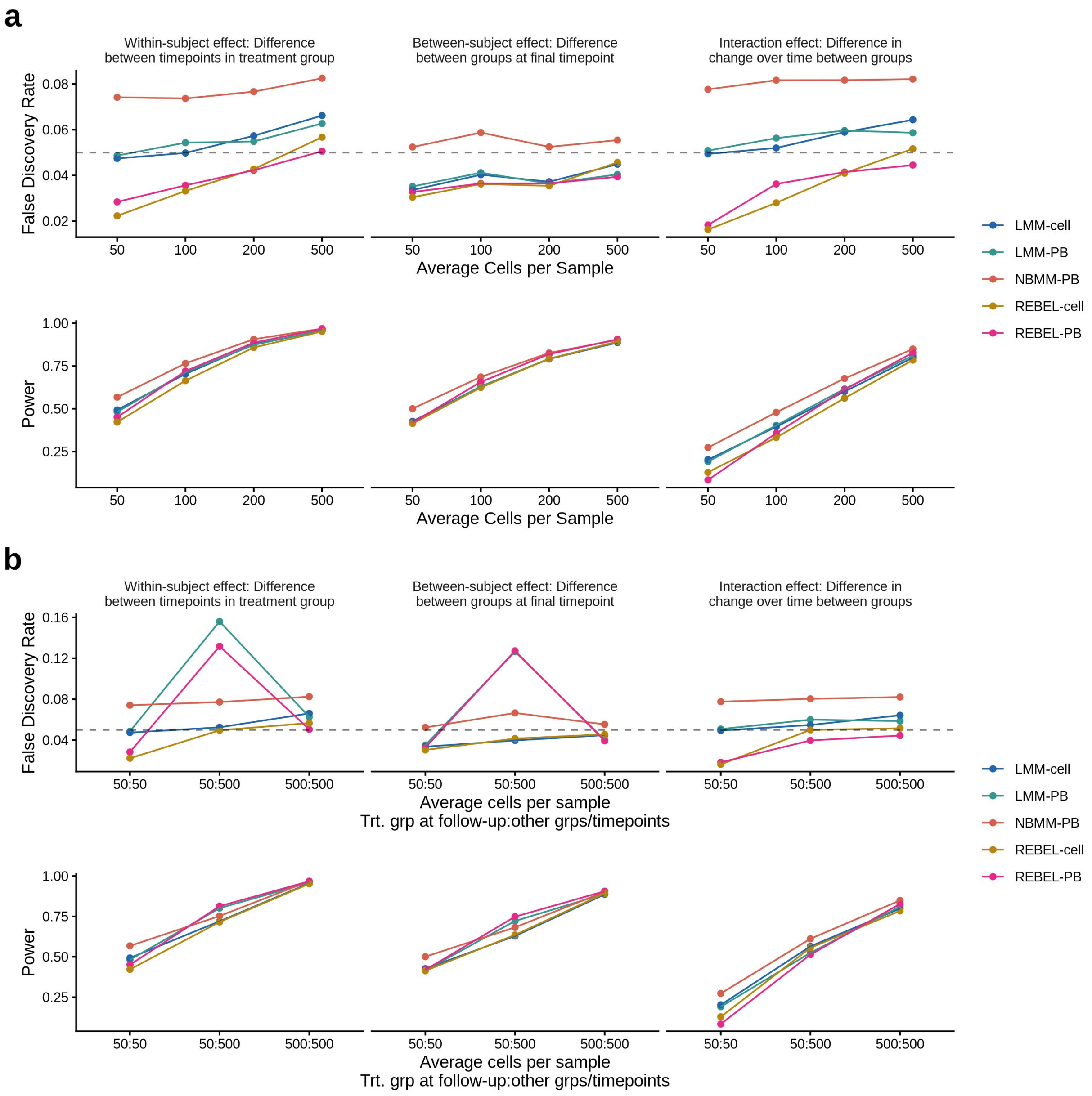


Figure S5: Observed false discovery rate (FDR) and power for traditional mixed-effects models and REBEL implementations under alternative cell-abundance scenarios. (a) Results for simulations in which the average number of cells per sample was varied uniformly across all groups and timepoints. (b) Results comparing balanced and imbalanced cell abundance scenarios with the x-axis labels contain two numbers with the first number showing the average number of cells in the treatment group at the final timepoint and the second number representing the average number of cells for samples in all other group/timepoint combinations. Results are shown for the three contrasts of interest. Statistical significance was defined using a Benjamini-Hochberg adjusted $p$-value threshold of 0.05. Dashed lines in the FDR plots denote the nominal 5% false discovery rate.

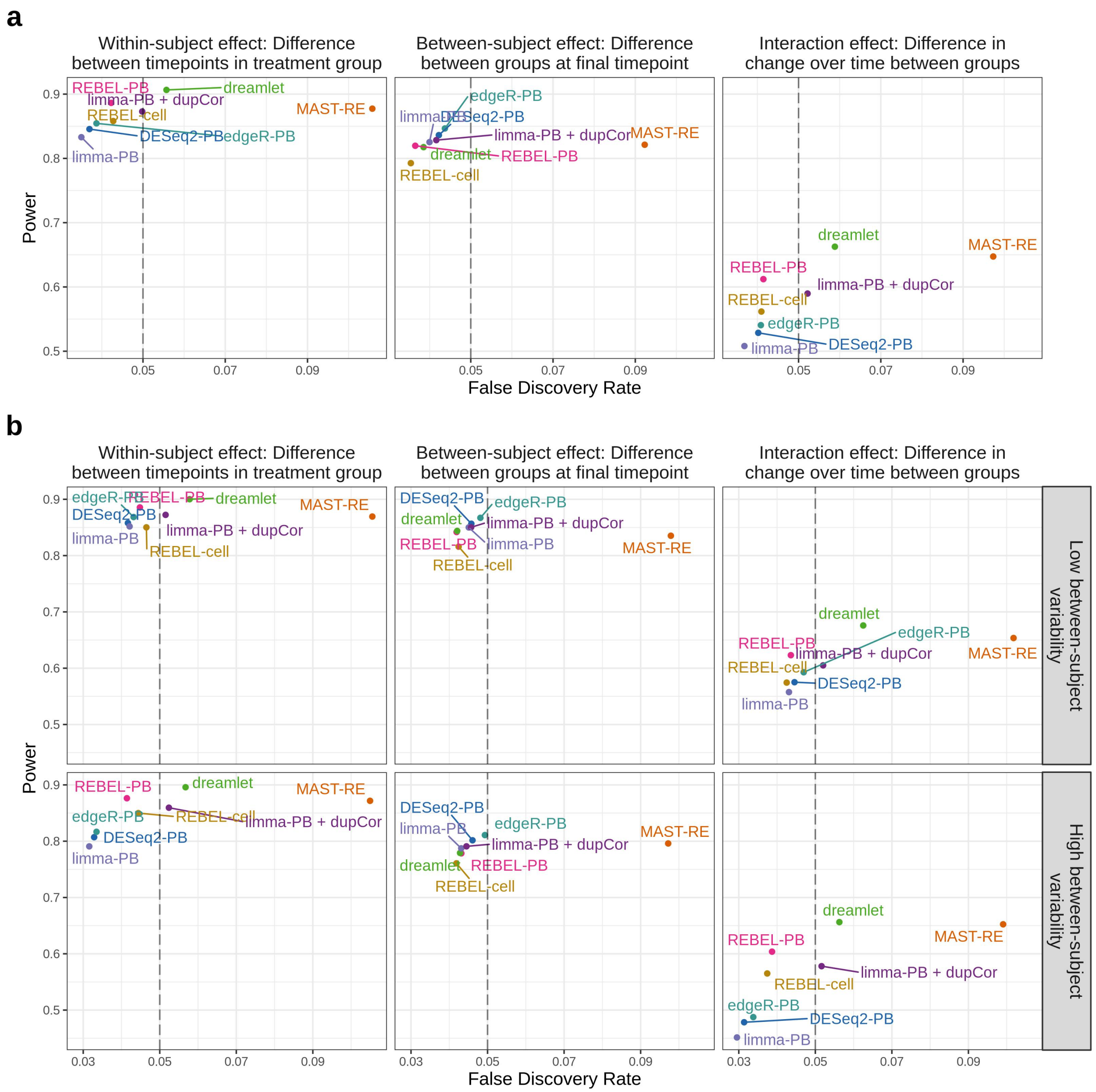


Figure S6: Observed false discovery rate (FDR) versus power for RNA-seq methods that account for correlation, including DESeq2-PB and edgeR-PB, under (a) baseline simulation settings and b) high and low between-subject variability settings. Ten simulated datasets were analyzed for each scenario and the FDR and power were averaged across simulation replicates. Results are shown across the three tests of interest. Results are shown for the three contrasts of interest. Statistical significance was defined using a Benjamini-Hochberg adjusted $p$-value threshold of 0.05. Dashed lines denote the nominal 5% false discovery rate.

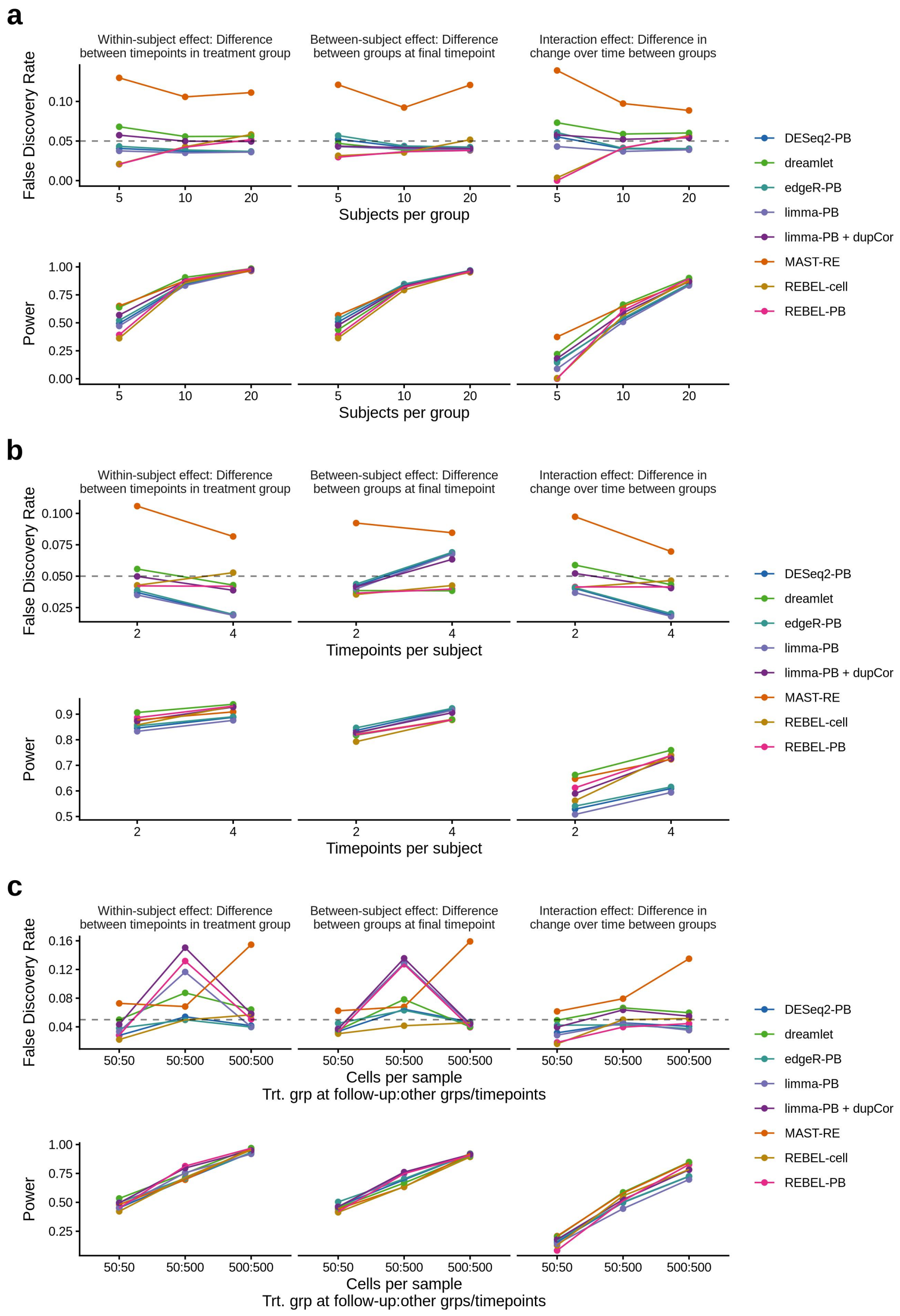


Figure S7: Observed false discovery rate (FDR) and power for RNA-seq methods that account for correlation, including DESeq2-PB and edgeR-PB, across different scenarios of (a) number of subjects per group, (b) Number of timepoints, and (c) average number of cells per group. In (c) the x-axis labels contain two numbers with the first number showing the average number of cells in the treatment group at the final timepoint and the second number representing the average number of cells for samples in all other group/timepoint combinations. Results are shown across the three tests of interest. Ten simulated datasets were analyzed for each scenario and the FDR and power were averaged across datasets. A 0.05 threshold was used to determine significance. The dashed lines in the FDR plots represent the nominal rate of 5% false discoveries.

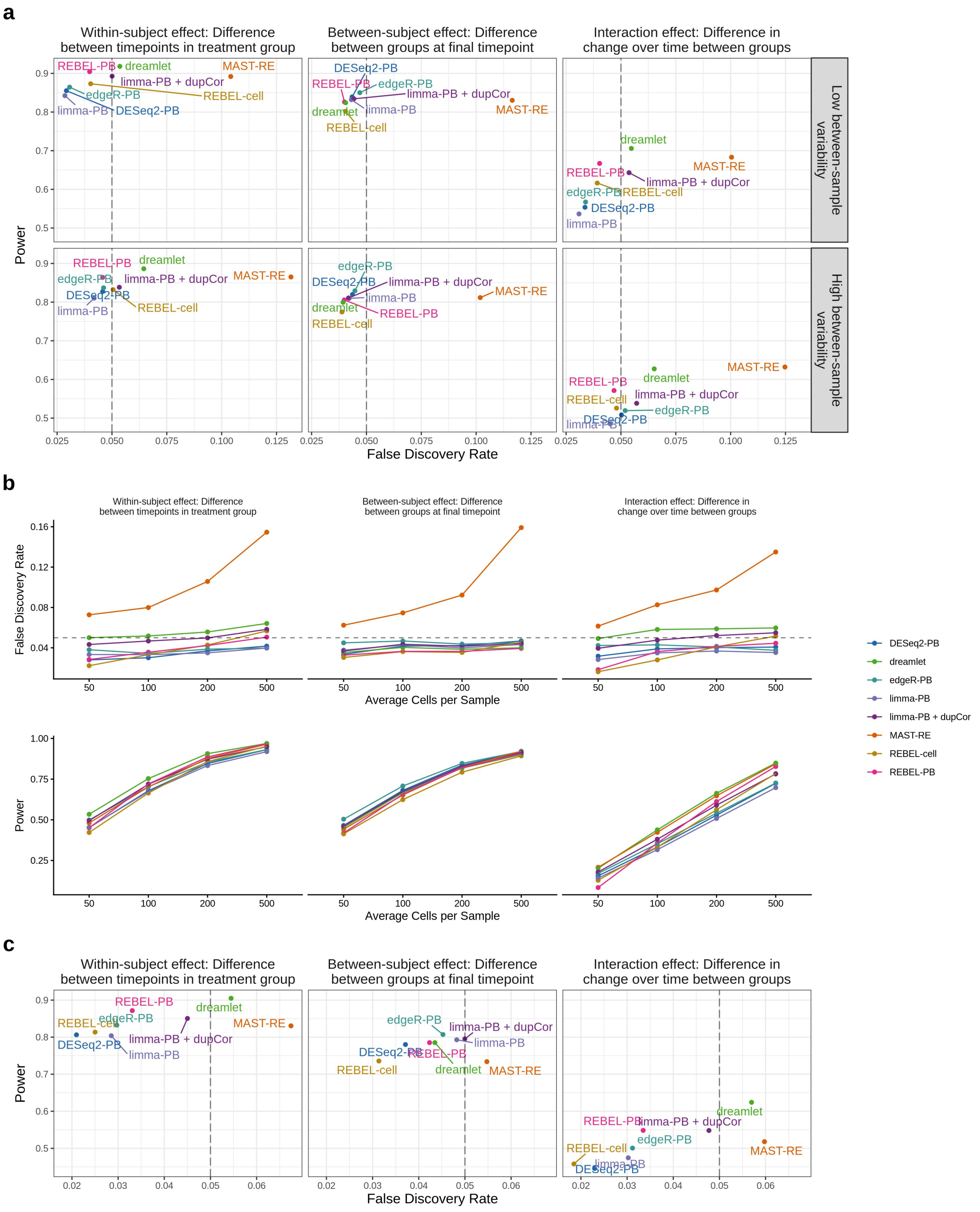


Figure S8: Observed false discovery rate (FDR) and power for RNA-seq methods that account for correlation, including DESeq2-PB and edgeR-PB, across alternative simulation scenarios. (a) Observed FDR versus power under high and low between-sample variability settings. (b) Observed FDR and power across simulations with differing average numbers of cells per sample. (c) Observed FDR versus power for the CD4$^+$ T-cell simulation scenario. Ten simulated datasets were analyzed for each scenario, and FDR and power were averaged across simulation replicates. Results are shown for the three contrasts of interest. Statistical significance was defined using a Benjamini–Hochberg adjusted $p$-value threshold of 0.05. Dashed lines in the FDR panels denote the nominal 5% false discovery rate.

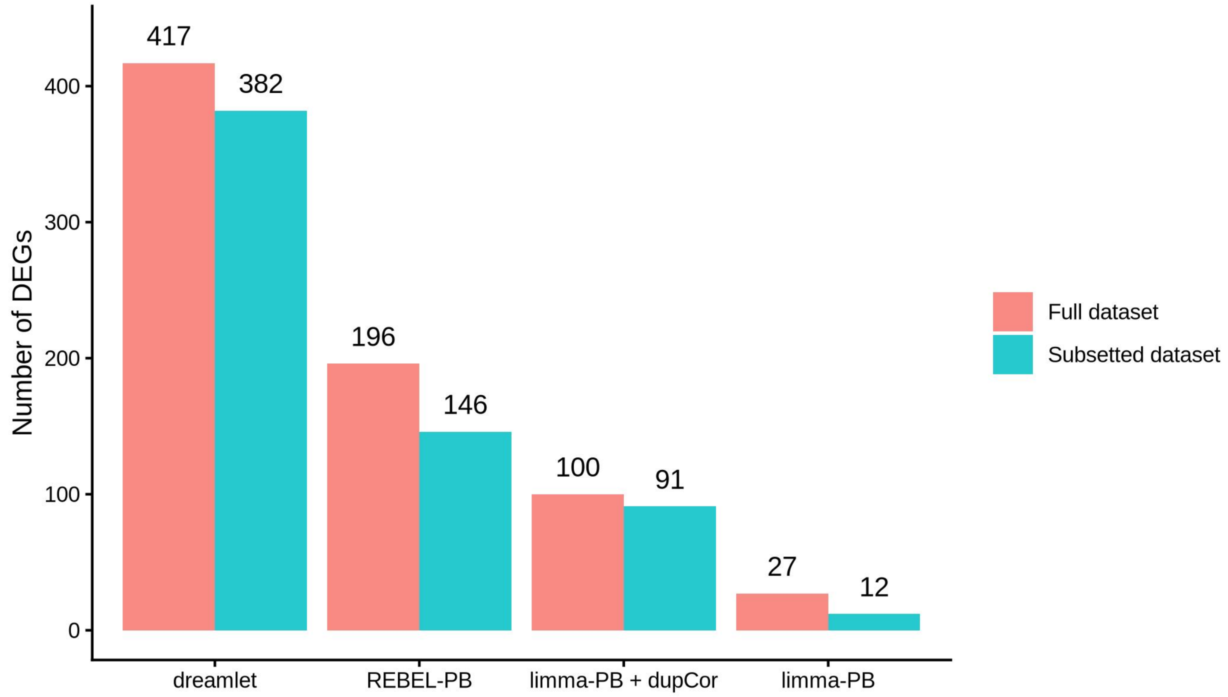


Figure S9: The number of statistically significant DEGs identified by each method using the full dataset (pink) and the downsampled dataset (blue, maximum 500 cells per sample) are shown. Methods are restricted to those applied to both datasets.
